# Brain Network Differences in Second Language Learning Depend on Individual Competencies

**DOI:** 10.1101/2025.09.28.679014

**Authors:** Nicole H. Skieresz, Sandy C. Marca, Micah M. Murray, Thomas P. Reber, Nicolas Rothen

**Author notes:** These authors contributed equally to this work (Micah M. Murray, Thomas P. Reber, Nicolas Rothen). Corresponding Author: Nicole H. Skieresz.

## Abstract

Integrating new words into an existing semantic network is a core challenge of second language (L2) acquisition. We investigated how evidence-based learning strategies and individual performance shape the neurocognitive dynamics of vocabulary learning. Eighty-three adults with German or French as their native language (L1) learned 48 Finnish (L2) nouns over 14 days using a mobile app that systematically varied retrieval practice, corrective feedback, multisensory learning, and distributed learning. Before and after training, EEG was recorded during a translation recognition task designed to elicit the N400, an index of semantic integration. Vocabulary accuracy increased from 0.41% pre-learning to 75.5% post- learning (*d_z_* = 3.96), and the N400 incongruity effect increased significantly, *F*(1, 75) = 99.52, *p* < .001, *η²_g_* = .32, reflecting successful integration of new L2 words into the mental lexicon. High performers showed larger N400 responses and distinct ERP template-map preponderance (i.e., the proportion of epoch time points assigned to a given template map) indicating more efficient and specialized neural processing. Despite systematic manipulation of learning strategies, no single approach yielded consistent behavioral or neural advantages, suggesting that overall exposure and cumulative practice—rather than any specific strategy— were the key drivers of robust learning. ERP template-map analyses further revealed that learning not only amplified neural responses but also shifted the preponderance of maps in the N400 window, signaling a qualitative reorganization of semantic processing. These findings bridge cognitive neuroscience and language education, suggesting that the depth and success of vocabulary learning may depend more on the degree of integration achieved than on the specific instructional strategy employed.

## 1 Introduction

Language comprehension depends on the rapid integration of novel lexical items into an existing semantic network. The N400 component—an event-related potential (ERP) that typically peaks around 400 ms post-stimulus—is widely recognized for its sensitivity to semantic congruency. Words incongruent with the preceding context elicit larger (more negative) N400 amplitudes. This phenomenon, known as the N400 incongruity effect, is interpreted as reflecting increased semantic processing demands (Kutas & Hillyard, 1980).

The scalp distribution of the N400 is dynamic, varying with stimulus type and learning stage. In standard visual-word paradigms, the N400 emerges as a monophasic negativity between 200 and 600 ms, with maximal amplitude over centro-parietal electrodes and a slight right-hemisphere bias (Kutas & Federmeier, 2011). For isolated written words, the N400 typically peaks at centro-parietal sites, while more frontal distributions are observed during early learning or in highly contextual scenarios (Van Petten & Luka, 2012). In a recent ERP study of novel-word learning, Armstrong et al. (2024) traced picture-word associative learning from initial encoding to multi-day retention and observed this topographic shift: early learning elicited a predominantly fronto-central N400, which later shifted toward the canonical centro-parietal maximum as representations consolidated.

Crucially, the N400 effect also emerges in translation paradigms, where participants view word pairs in two languages: correctly matched (“congruent”) translation pairs elicit smaller N400 amplitudes than mismatched (“incongruent”) pairs (Midgley et al., 2009; Pu et al., 2016; Yum et al., 2014), reflecting sensitivity to semantic mismatch. This property has established the N400 as a key neural marker in second language (L2) vocabulary acquisition research. Pu et al. (2016) demonstrated that learners initially display no reliable N400 incongruity effect between congruent and incongruent L2–L1 translation pairs, but following vocabulary training, a robust N400 incongruity effect emerges, indicating successful integration of new L2 words into semantic memory. Similarly, Yum et al. (2014) found that rapid learners exhibited increased N400 amplitudes to incorrect translations early in training.

Recent studies in adult learners further underscore the N400’s sensitivity to lexical-semantic learning. McLaughlin et al. (2004) demonstrated that an N400 incongruity effect can emerge after only minimal L2 exposure, indicating rapid semantic integration. Elgort et al. (2014) further showed that L2 proficiency influences both behavioral performance and N400 topography: high-proficiency learners exhibited stronger semantic-relatedness effects with a canonical centro-parietal distribution, whereas low-proficiency learners displayed attenuated amplitudes and delayed peaks. Qi et al. (2017) reported that pre-training N400 amplitudes to native language stimuli predicted subsequent vocabulary and grammar learning in an artificial language. García-Gámez and Macizo (2022) demonstrated that semantic training involving word-picture pairs and categorization tasks elicited stronger N400 incongruity effects than lexical training with L1–L2 word associations, suggesting deeper semantic encoding. Together, these findings support the N400 as a neural marker of individual learning potential, highlighting that proficiency not only predicts learning outcomes but also modulates the component’s amplitude, timing, and scalp distribution.

Beyond semantic congruency, N400 amplitude also reflects lexical-access fluency (Van Petten & Luka, 2012) and, within predictive-coding frameworks, indexes prediction error (Lau et al., 2008; Kuperberg & Jaeger, 2016). Reduced N400 amplitudes for contextually expected stimuli are thought to reflect minimized discrepancies between predicted and encountered semantic information. Importantly, N400 modulations extend to multisensory stimuli (Kutas & Federmeier, 2011) and have even been observed in non-human species such as dogs (Boros et al., 2024; Murray et al., 2024), highlighting its evolutionary robustness as a marker of semantic processing.

Despite a growing body of ERP research in L2 acquisition, few studies have directly examined how specific learning strategies affect neural indices of vocabulary acquisition (e.g., García-Gámez & Macizo, 2022; Li et al., 2023; Soskey et al., 2016). The present study addresses this gap by combining the ERP translation recognition paradigm from Pu et al. (2016) with a learning intervention incorporating four evidence-based strategies. These strategies were selected based on empirically supported principles outlined by Reber and Rothen (2018), which have been identified as particularly effective for promoting retention in digital vocabulary learning: retrieval practice, corrective feedback, multisensory encoding, and distributed learning.

Retrieval practice involves actively recalling learned items rather than passively re-exposing learners to them. This strategy has been shown to enhance long-term retention by reinforcing memory traces through active recall (Roediger & Karpicke, 2006). In the context of vocabulary learning, retrieval is associated with more stable lexical associations and improved recall accuracy (Barcroft, 2007; Kang et al., 2013). It is hypothesized that retrieval practice sharpens lexical access and supports more robust semantic integration, which may be reflected in larger N400 incongruity effects after learning—that is, a greater neural differentiation between congruent and incongruent word pairs. However, some studies report reduced N400 amplitudes following successful retrieval, potentially reflecting more automatic or efficient semantic access (Li et al., 2023). These findings suggest that N400 modulation may vary with learning stage, task demands, or the strength of lexical-semantic associations. While behavioral benefits of retrieval practice are well established, neurophysiological evidence remains limited, and further research is needed to clarify how such strategies modulate ERP components during vocabulary acquisition.

Corrective feedback during learning is provided by immediately presenting the correct information after a failed or erroneous retrieval attempt. This approach supports the formation of accurate internal representations of word meanings by directly addressing incorrect associations (Nicol & Macfarlane-Dick, 2006; Pashler et al., 2005). It is hypothesized that this process supports the development of more distinct lexical-semantic mappings, which may lead to greater neural differentiation between congruent and incongruent word pairs. However, while its pedagogical benefits are well established, ERP-based research specifically investigating the neural correlates of feedback in vocabulary learning remains sparse.

Multisensory learning refers to instructional conditions in which information is presented through more than one sensory modality, such as visual, auditory, or kinesthetic channels. Multisensory inputs have been shown to improve recall and support deeper semantic encoding (Shams & Seitz, 2008; Thelen & Murray, 2013). Empirical studies demonstrate that multisensory, compared to unisensory, inputs lead to more robust and differentiated neural representations, including stronger and earlier ERP components (Murray et al., 2004) as well as enhanced free recall performance following audiovisual compared to auditory-only encoding (Atkin et al., 2023; Soto-Faraco & Spence, 2025).

Distributed learning refers to study conditions in which study sessions are spaced over time rather than amassed within a single block. This spacing enables memory traces to consolidate between sessions and has been shown to enhance long-term retention and semantic stability (Cepeda et al., 2006; Edmonds et al., 2021). Neuroimaging research indicates that spaced learning strengthens lexical-semantic representations by increasing neural pattern similarity across repetitions, suggesting more consistent and consolidated memory traces (Feng et al., 2019). It is hypothesized that this consolidation-driven strengthening supports the formation of more stable semantic expectations, potentially resulting in more robust N400 incongruity effects, although this link remains to be clarified.

To test the impact of these four evidence-based learning strategies—retrieval practice, corrective feedback, multisensory encoding, and distributed learning—on neural markers of vocabulary learning, we employed a translation recognition ERP paradigm and analyzed the data using both canonical ERP methods and an electrical neuroimaging framework based on global measures of the electric field at the scalp (Murray et al., 2008) to characterize the spatiotemporal dynamics of the N400. We further explored how individual learning success (high vs. low performers) modulates these strategy effects.

Guided by prior ERP research on L2 vocabulary acquisition (e.g., Pu et al., 2016; García-Gámez & Macizo, 2022) and learning theory from cognitive psychology (e.g., Roediger & Karpicke, 2006; Cepeda et al., 2006), we formulated four hypotheses:

- Hypothesis 1 (Replication): Successful L2 vocabulary learning will elicit a robust N400 incongruity effect from pre- to post-test, replicating prior findings of semantic integration.
- Hypothesis 2 (Neural Effects of Strategy): Learning strategies that promote lexical-semantic integration—namely retrieval practice, corrective feedback, multisensory encoding, and distributed learning—will enhance the N400 incongruity effect after training.
- Hypothesis 3 (Behavioral Impact): Items learned under these strategies will show superior learning outcomes compared to items trained without them.
- Hypothesis 4 (Individual Differences): High-performing learners will show larger N400 incongruity effects than low performers, indicating more efficient and robust lexical-semantic integration.

Together, these hypotheses aim to clarify how evidence-based learning strategies modulate both the neural and behavioral outcomes of L2 vocabulary learning.

## 2 Methods

### 2.1 Participants

Eighty-three healthy adults (M_age_ = 28.59 years; range = 18–40; 54 females) with normal or corrected-to-normal vision participated as part of a larger project. Participants were either native German or French speakers or had advanced proficiency in one of these languages (N_German_ = 50; N_French_ = 33). Recruitment was conducted via the UniDistance Suisse online participant database. All participants provided written informed consent and completed a comprehensive cognitive test battery. Six individuals were excluded from EEG analysis due to incomplete neurophysiological assessment or poor data quality. Participants received either compensation for up to eleven experimental hours or a payment of 200 CHF upon study completion. The ethics committee of UniDistance Suisse approved all procedures (Nr. 2019-12-00002).

### 2.2 Design

The study employed a mixed factorial design incorporating both within-subject factors—Session (pre vs. post); Congruency (congruent vs. incongruent); Retrieval Practice (high vs. low); Corrective Feedback (corrective vs. non-corrective); Multisensory Learning at Encoding (multisensory vs. unisensory); Multisensory Learning at Retrieval (multisensory vs. unisensory)—and two between-subject factors—Performance Group (high vs. low performers) and Distributed Learning (distributed vs. massed), implemented across three sequential phases: pre-learning assessment, vocabulary learning intervention, and post-learning reassessment.

Dependent variables included vocabulary-test accuracy (percent correct) and translation recognition task response-bias measures (*d′*, *c*, ln *β*). Neurophysiological measures comprised mean N400 amplitude (300–500 ms, six channels), preponderance of ERP template maps (topographic clustering), Global Field Power (GFP), and Global Map Dissimilarity (GMD).

### 2.3 Materials

#### 2.3.1 Stimuli

Forty-eight Finnish nouns (L2) were each paired with a German or French translation (L1) and consistently presented across phases. Language direction (L1–L2 vs. L2–L1) was varied across the vocabulary test, the translation recognition task, and the mobile-app intervention, but was held constant within each participant across phases. Across the sample, items were intended to appear approximately equally often in L1–L2 and L2–L1 (50:50). Due to a small number of stimulus exclusions (see below and *Supplementary Table S1*), the final analyzed dataset deviates slightly from the intended balance. All stimuli were presented in three modalities during the intervention: auditory, visual, and pictorial. Auditory stimuli were sourced from Universal-Soundbank.com, freesound.org, or generated via Google Translate (a female voice). Visual stimuli (written form) were presented in dark blue on a white background, with associated images, adhering to a 4:3 aspect ratio, sourced mainly from Wikimedia Commons or created using Microsoft Paint 3D. Stimulus characteristics are detailed in *Table 1*, with the complete list provided in *Supplementary Table S1*.

**Table 1.** Description of the Stimuli by Language.

| Language | Number of syllables |  | Word length |  | Zipf Frequency |  |
| --- | --- | --- | --- | --- | --- | --- |
|  | M | SD | M | SD | M | SD |
| German | 1.72 | 0.54 | 5.04 | 1.23 | 4.02 | 0.56 |
| French | 2 | 0.51 | 5.64 | 1.24 | 4.10 | 0.56 |
| Finnish | 2.26 | 0.44 | 5.06 | 0.99 | 3.91 | 0.76 |

*Note.* The Zipf frequency of a word is the base-10 logarithm of the number of times it appears per billion words. Frequencies were determined using the wordfreq package (Speer, 2022).

Four items contained typographical errors. Three of these affected all French-speaking participants and were excluded from all analyses. In a fourth case, the Finnish word *huili* (instead of *huilu*, “flute”) was presented during the intervention phase for four German-speaking participants, while *huilu* was used in the test phases. Due to this inconsistency, data for this specific word were excluded for these participants. Items with typographical errors are marked with an asterisk (*) in *Supplementary Table S1*.

Additionally, nine Finnish words used in the experiment did not correspond exactly to the intended German source words due to translation errors during stimulus preparation (marked with a dagger [^†^] in the Supplement). These translation discrepancies are detailed in the Supplementary Table and Table Note. All translation errors were present in the stimuli as presented to participants. Since none of the participants had prior knowledge of Finnish, these errors are not expected to have affected task performance or interpretation of results.

#### 2.3.2 Vocabulary Test and Scoring

The vocabulary test was administered as part of a cognitive test battery, involving cued recall tasks where participants translated between L1 and L2. Stimuli appeared centrally (black on grey background) via a standard 14-inch Hewlett-Packard laptop.

For each participant and test session, accuracy was calculated as the proportion of correctly translated items. Responses were scored using a semi-automated algorithm based on orthographic similarity (optimal string alignment; see van der Loo, 2014, for stringdist package), allowing for minor spelling errors and variations. A response was classified as correct if it matched the target translation exactly or differed by only a single character (“perfect” or “close” responses). All other responses were scored as incorrect. In cases where a response was equally close to more than one possible target word (“tie”), the item was excluded from scoring to ensure unambiguous classification.

Additional cognitive tests included: Adapted French l’EVIP (Dunn & Thériault- Whalen, 1993) to assess L1 vocabulary size, Test of Variables of Attention (TOVA; Leark et al., 2008) for sustained attention, Standard Progressive Matrices (Raven et al., 1958) measuring fluid intelligence, a serial order reconstruction task (Leclercq & Majerus, 2010; Gorin & Majerus, 2019; Majerus et al., 2006) for short-term order memory, a non-word short-term recognition task (Leclercq & Majerus, 2010; Gorin & Majerus, 2019; Majerus et al., 2006) assessing item-specific short-term memory, and a tapping continuation task (Dalla Bella et al., 2017; Gorin & Majerus, 2019) for rhythmic processing. Tests were programmed and conducted using Lab.js (version 20.2.4; Henninger et al., 2020).

#### 2.3.3 Translation Recognition Task

A translation recognition task, adapted from Pu et al. (2016), was used to assess the N400 incongruity effect. The task was conducted in Octave (version 4.0.3) with PsychToolbox extensions (3.0.14) on a Debian system.

In each trial, two words were presented in black color sequentially at the center of a 24-inch LCD color monitor (HP LA2405x) with a light grey background. Each word appeared for 800 ms, with no interstimulus interval. The intertrial interval was fixed at 1,000 ms and included a jitter of 300–600 ms.

Participants used the left or right arrow key to indicate whether they believed the second word was a correct translation of the first word. Immediately afterward, they responded to a second prompt by again using the arrow keys to indicate whether their decision was based on knowledge or guessing. Task duration ranged from approximately 30 to 50 minutes with optional breaks between blocks.

Each of the 48 stimuli appeared once in a correct (congruent) and once in an incorrect (incongruent) translation per block, yielding 96 trials per block. Across four blocks—with alternating translation directions—participants completed either two blocks of L2–L1 followed by two blocks of L1–L2, or vice versa (two blocks L1–L2 followed by two blocks L2–L1). The order of these four blocks was counterbalanced across participants. Each block comprised 96 trials, with each of the 48 stimuli appearing once in a congruent and once in an incongruent pairing, yielding a total of 384 experimental trials per session.

Incongruent pairs were generated by randomly combining words from the same stimulus set, ensuring semantic mismatch. In the first version of the task (N = 16), incongruent combinations were newly generated for each block, such that each incongruent pairing appeared at least once per session. This design introduced a potential learning artifact, as participants could begin to infer incongruent pairs through repetition. To address this, a second version (N = 67) fixed all incongruent combinations across blocks and sessions. In both versions, congruent pairs remained constant. Response key assignments were randomized and counterbalanced in the first version but fixed in the second version for consistency across sessions.

### 2.4 Procedure

All sessions were conducted at the same UniDistance Suisse site for each participant—either one of the German-speaking locations (Naters or Brig) or the French-speaking site (Sierre). The study comprised a pre-learning phase (Sessions 1 and 2), a 14-day intervention phase, and a post-learning phase (Session 3). *Figure 1* shows a schematic overview of the procedure.

**Figure 1.**
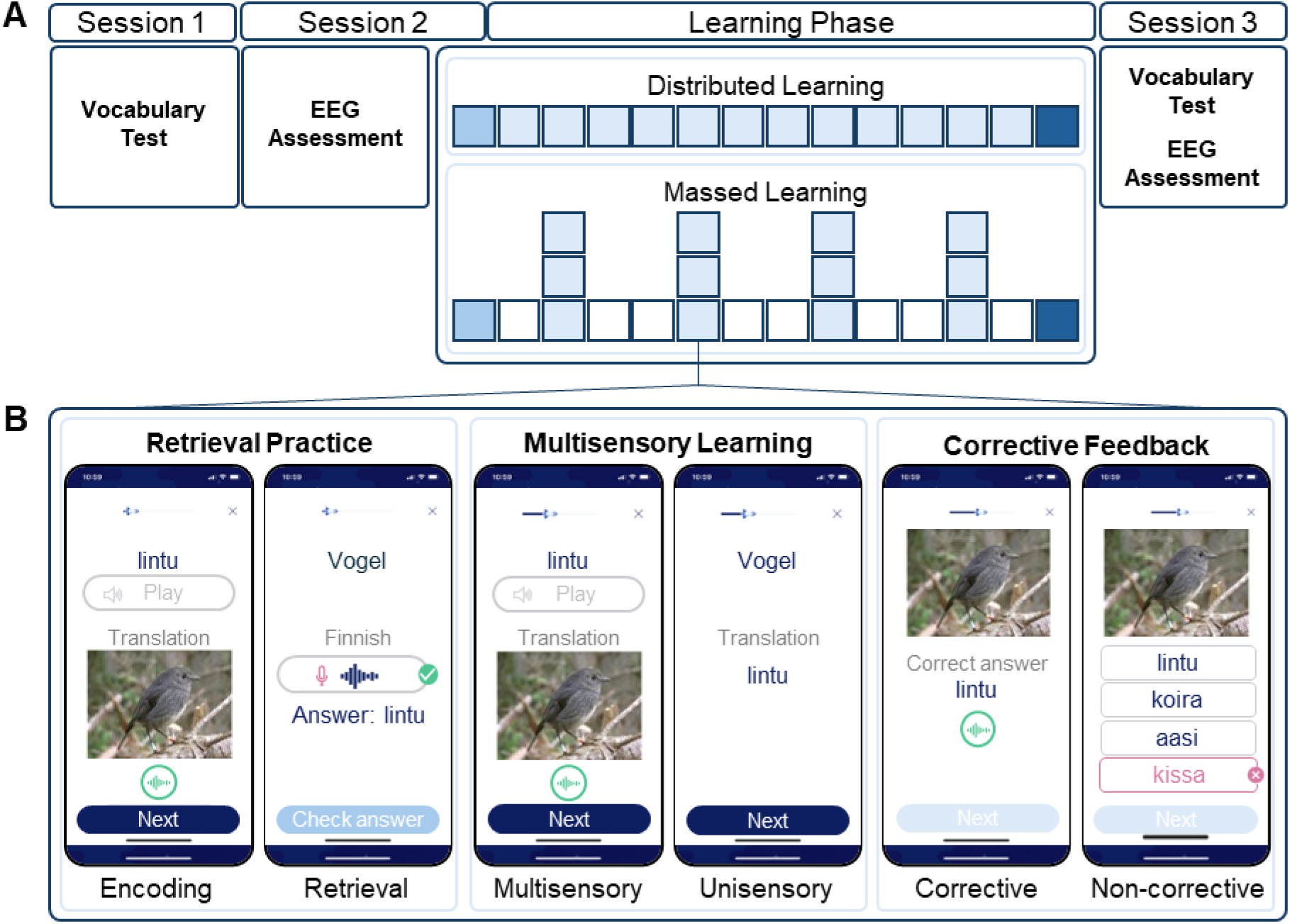
Overview of the Experimental Procedure and Implementation of Learning Strategies. *Note.* (A) Experimental phases: pre-learning (Session 1: baseline vocabulary testing; Session 2: EEG assessment and initiation of the 14-day mobile-app intervention; blue = learning days; white = rest days); and post-learning (Session 3: vocabulary testing and EEG reassessment). Participants were randomized to distributed or massed schedules. (B) Example of word presentation in the mobile app during intervention. Interface elements (e.g., buttons and labels) are shown in English for illustration; the actual app was presented in German or French (L1) and Finnish (L2). Stimuli were allocated by retrieval proportion (70% retrieval practice; 30% restudy), modality (50% multisensory; 50% unisensory), and feedback type (50% corrective; 50% non-corrective). The intervention phase comprised 48 initial encoding trials, 12 daily sessions of 48 trials each, and a final retrieval session of 20 trials (total = 644 trials per participant).

Session 1 (approximately 2.5 hours) involved cognitive and vocabulary testing and was conducted individually or in small groups (51 individually, 22 pairs, 10 groups of three).

Session 2 (approximately 2 hours) consisted of a demographic questionnaire, visual acuity testing using Landolt rings (optikschweiz.ch), and EEG setup. Participants were seated approximately 60 cm from the computer screen, and a chin rest was used to minimize head movements and maintain constant viewing distance. The neurophysiological assessment began with a 3-minute resting-state EEG recording (eyes open), followed by two EEG tasks—a translation recognition and a fast periodic visual stimulation (FPVS) oddball paradigm—administered in random order. Only the translation recognition task is reported in the present manuscript. The EEG session concluded with detailed instructions for the mobile app-based vocabulary training.

Session 3 (approximately 2 hours) comprised a repeated vocabulary test and EEG assessments. Six participants completed the EEG tasks before the vocabulary test to minimize order effects. The order of the two EEG tasks was kept consistent with Session 2.

### 2.5 Electrophysiological Recording and Pre-processing

EEG was recorded with a 64-channel sponge-based electrode system (RNet; Brain Products GmbH), arranged according to the extended international 10–20 system (ground: FPz, reference: Cz), with impedances kept below 30 kΩ. Signals were digitized at 1,000 Hz using a BrainAmp amplifier (Brain Products GmbH) and recorded on a Lenovo laptop (Windows 10 Home, 8 GB RAM).

Pre-processing was performed using EEGLAB (version 2024.1; Delorme & Makeig, 2004; The MathWorks, Inc., 2022) and ERPLAB (version 12.00; Lopez-Calderon & Luck, 2014) and comprised downsampling to 250 Hz; band-pass filtering between 0.1 and 35 Hz; continuous artifact rejection; interpolation of broken or artifact-contaminated channels; re-referencing to the common average; peri-stimulus epoching (−200–800 ms); pre-stimulus baseline correction (−200–0 ms); artifact rejection on epochs (±150 µV), and removal of eye- blink artifacts. Following pre-processing, an average of 302 ± 68 epochs per participant pre-learning and 305 ± 64 epochs post-learning were retained.

### 2.6 Data Analysis

#### 2.6.1 Statistical Analyses & Variables

T-tests and mixed ANOVAs were conducted in RStudio (version 2024.12.0) with *p* = .05 as the threshold for statistical significance.

Behavioral dependent variables included vocabulary test accuracy (percent correct) and response bias indices from the translation recognition task (sensitivity (*d*’), criterion location (*c*), likelihood ratio (ln *β*)). Response-bias indices were calculated as per Macmillan & Creelman (2005), with hit and false alarm rates corrected, using Snodgrass & Corwin’s (1988) formula (adding 0.5 and dividing by 𝑁 + 1, where 𝑁 is the number of congruent or incongruent trials).

The specific computations were:

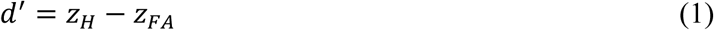

The criterion location was computed as

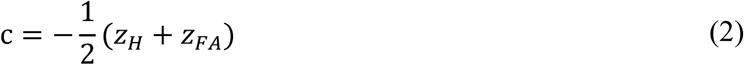

The likelihood ratio was computed as

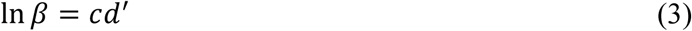

where 𝑧_𝐻_ and 𝑧_𝐹𝐴_ denote the z-transformed hit rate and false alarm rate, respectively.

For further group-level analyses, a median split was performed on vocabulary gain (i.e., post-test minus pre-test accuracy) across all participants; those with gains equal to or above the sample median were classified as “high performers,” and those below as “low performers.”

Electrophysiological dependent variables encompassed mean N400 amplitudes measured at six centro-parietal electrodes (C3, Cz, C4, P3, Pz, P4), reflecting the canonical centro-parietal distribution of the mature N400 (Kutas & Federmeier, 2011; Van Petten & Luka, 2012). We quantified mean amplitudes in the 300–500 ms window post-stimulus onset (Šoškić et al., 2021). To capture global spatiotemporal dynamics, we performed topographic clustering and derived template-map preponderance (proportion of time frames assigned to each map within predefined epochs; time frames = TF) and used RAGU to compute Global Field Power (GFP) and Global Map Dissimilarity (GMD). Effects in GFP/GMD were assessed with the permutation-based global pAUC statistic (5,000 permutations). In the Results, we report the permutation *p* values associated with the global pAUC tests (formatted as “global pAUC, *p* = …”).

#### 2.6.2 Topographic Clustering, Global Field Power, and Global Map Dissimilarity

Analyses of neural activation were conducted using the freeware Cartool (version 5.02; Brunet et al., 2011) for topographic clustering and freeware RAGU (compiled 24 November 2020; Koenig et al., 2011) for GFP and GMD.

Topographic clustering decomposed continuous ERPs into transient but stable ERP template maps that reflect discrete functional brain states. Grand-averaged ERPs per condition (pre-congruent, pre-incongruent, post-congruent, post-incongruent) were imported into Cartool and spatially filtered to enhance signal quality. Segmentation was performed using the Topographic Atomize and Agglomerate Hierarchical Clustering (T-AAHC) algorithm with a minimum segment length of 10 time frames (TF). The resulting group-level ERP template maps were then backfitted to individual ERPs via spatial correlation to derive map labels and preponderance.

GFP analysis assessed differences in the strength of neural responses across conditions. GFP reflects the overall amplitude of electric brain activity, independent of topographic configuration. At each time frame 𝑡, GFP was computed as the root mean square across the 𝑁 electrodes (Equation 4). Significance for GFP was evaluated within RAGU’s global pAUC framework using nonparametric permutation tests (5,000 permutations).

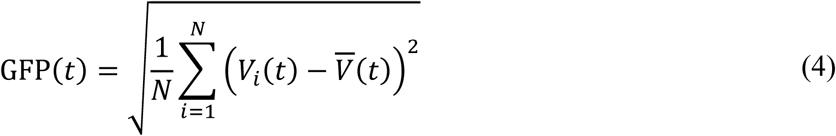

GMD analysis evaluated condition-related differences in scalp topographies for each pair of the four experimental conditions (pre-congruent, pre-incongruent, post-congruent, post-incongruent) across the full post-stimulus period (0–796 ms). In contrast to topographic clustering, which identifies recurring ERP template maps within predefined epochs, GMD provides a time-resolved measure of topographic dissimilarity between two conditions. For any two conditions 𝛼 and 𝛽, GMD at time 𝑡 was computed as:

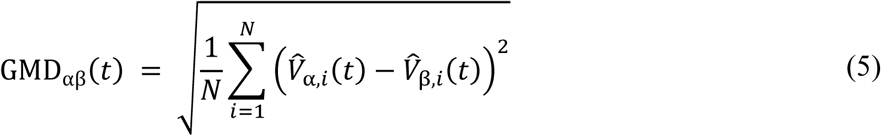

where 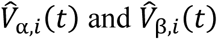 are the GFP-normalized potentials at electrode 𝑖 for conditions 𝛼 and 𝛽, respectively and 𝑁 is the number of electrodes. Statistical significance for GMD was likewise assessed via nonparametric permutation testing (5,000 permutations) within the global pAUC framework; in the Results we report the associated permutation *p* values.

## 3 Results

**3.1 Vocabulary Test**

Vocabulary test accuracy increased substantially from pre- to post-learning (see *Figure 2*). On average, participants scored *M* = 0.41% (*SD* = 1.19%; range 0–6.82%) before learning and *M* = 75.5% (*SD* = 19.1%; range 16.7–100%) after learning. A paired-samples t- test confirmed this change was highly significant, *t*(82) = 36.07, *p* < .001, *d_z_* = 3.96, 95% CI [3.32, 4.59], indicating a large learning effect.

**Figure 2.**
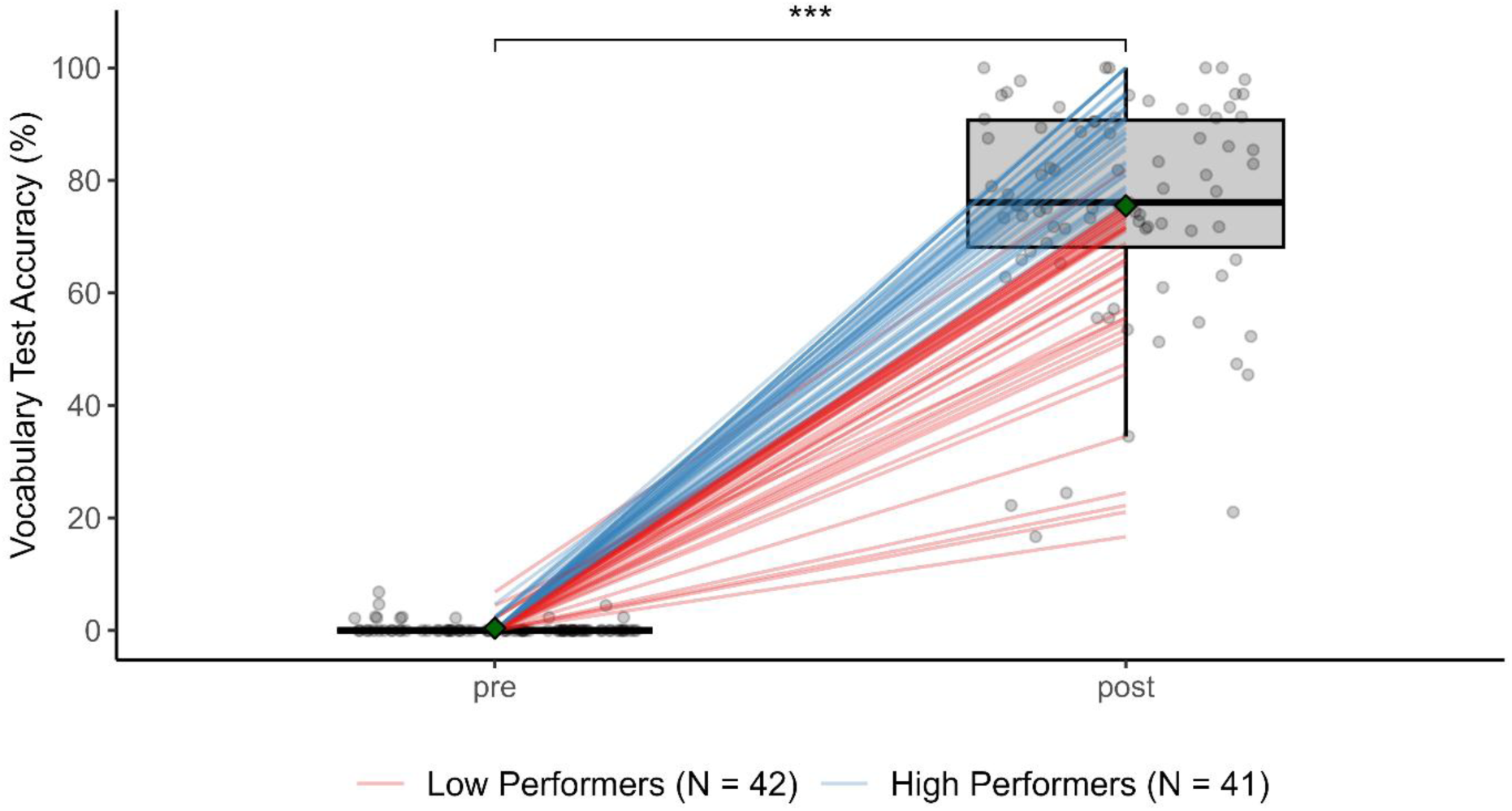
Vocabulary Test Accuracy Before and After Learning. *Note.* Boxplots show participants’ test accuracy (%) before and after learning. Gray circles are individual accuracy; colored lines connect paired measurements (low performers in red, high performers in blue). Green diamonds indicate group means, and asterisks denote significant within-group gains (*p* < .001).

None of the main effects of Retrieval Practice, Multisensory Learning at Encoding and Retrieval, or Distributed Learning were significant (all *p* > 0.116; see *Supplementary Table S2*). There was a small but significant main effect of Feedback Type, *F*(1, 80) = 10.95, *p* = 0.001, *η²_g_* = 0.03. However, follow-up analyses showed that this effect was driven by low performers: The improvement in accuracy (post–pre) was higher for items learned with corrective feedback (*M* = 61.8, *SD* = 22.6) than with non-corrective feedback (*M* = 52.8, *SD* = 17.1), *t*(41) = 2.96, *p* = .005, *d_z_* = 0.46, 95% CI [0.14, 0.77]. In contrast, among high performers, there was no significant difference in improvement in accuracy between corrective (*M* = 88.9, *SD* = 11.0) and non-corrective feedback (*M* = 85.7, *SD* = 13.7), *t*(39) = 1.58, *p* = .121, *d_z_* = 0.25, 95% CI [−0.07, 0.56].

### 3.2 Response-Bias Indices

Sensitivity (*d’*), Criterion *c* and likelihood ratio (ln *β*) were computed as described in the Methods. Sensitivity in the translation recognition task improved markedly from pre- to post-learning. The mean *d′* score increased from *M* = 0.59 (*SD* = 0.34) to *M* = 4.15 (*SD* = 1.01), indicating a substantial gain in participants’ ability to distinguish correct from incorrect translations. A repeated-measures ANOVA revealed significant main effects of Session, *F*(1, 75) = 1603.27, *p* < .001, *η²_g_* = .89, and Performance Group, *F*(1, 75) = 30.16, *p* < .001, *η²_g_* = .20 (all *p* > .05; see *Supplementary Table S3)*.

Criterion *c* value decreased from *M* = 0.28 (*SD* = 0.45) pre-learning to *M* = 0.12 (*SD* = 0.26) post-learning, indicating a shift toward less conservative responding. ANOVA results revealed significant main effects of Performance Group, *F*(1, 75) = 5.92, *p* = .017, *η²_g_* = .04 and Session, *F*(1, 75) = 9.49, *p* = .003, *η²_g_* = .05. The Session × Performance Group interaction was not significant, *F*(1, 75) = 0.75, *p* = .390, *η²_g_* = .00. No learning-strategy factors reached significance (all *p* > .26, *η²_g_* ≤ .01; see *Supplementary Table S4*), except for small group-specific interactions for Multisensory Learning at Retrieval (Performance Group × RM: *F*(1, 75) = 4.67, *p* = .034, *η²_g_* = .01) and Distributed Learning (Performance Group × DL: *F*(1, 73) = 5.40, *p* = .023, *η²_g_* = .07).

The mean ln *β* value increased from *M* = 0.15 (*SD* = 0.28) pre-learning to *M* = 0.43 (*SD* = 0.97) post-learning, indicating a tendency toward more conservative decisions post-learning—participants were less likely to identify a translation pair as correct without sufficient certainty. A significant main effect of Session was found, *F*(1, 75) = 6.75, *p* = .011, *η²_g_* = .04. No significant main effect of Performance Group, *F*(1, 75) = 3.63, *p* = .061, *η²_g_* = .03, and no significant Session × Performance Group interaction, *F*(1, 75) = 0.62, *p* = .432, *η²_g_* = .00, were observed. No learning strategy effects were significant (all *p* > .05).

### 3.3 N400 Incongruity Effect

To evaluate changes in semantic processing, we quantified the N400 incongruity effect as the amplitude difference between incongruent and congruent trials. A repeated-measures ANOVA revealed a significant main effect of Session, *F*(1, 75) = 99.52, *p* < .001, *η²_g_* = 0.32, indicating a larger incongruity effect post-learning. There was also a main effect of Performance Group, *F*(1, 75) = 14.44, *p* = .001, *η²_g_* = .11, with high performers showing greater N400 incongruity effects overall. Importantly, the Session × Performance Group interaction was significant, *F(*1, 75) = 6.79, *p* = .011, *η²_g_* = .03, reflecting that high performers exhibited a larger pre-to-post increase than low performers (see *Figure 3A*).

**Figure 3.**
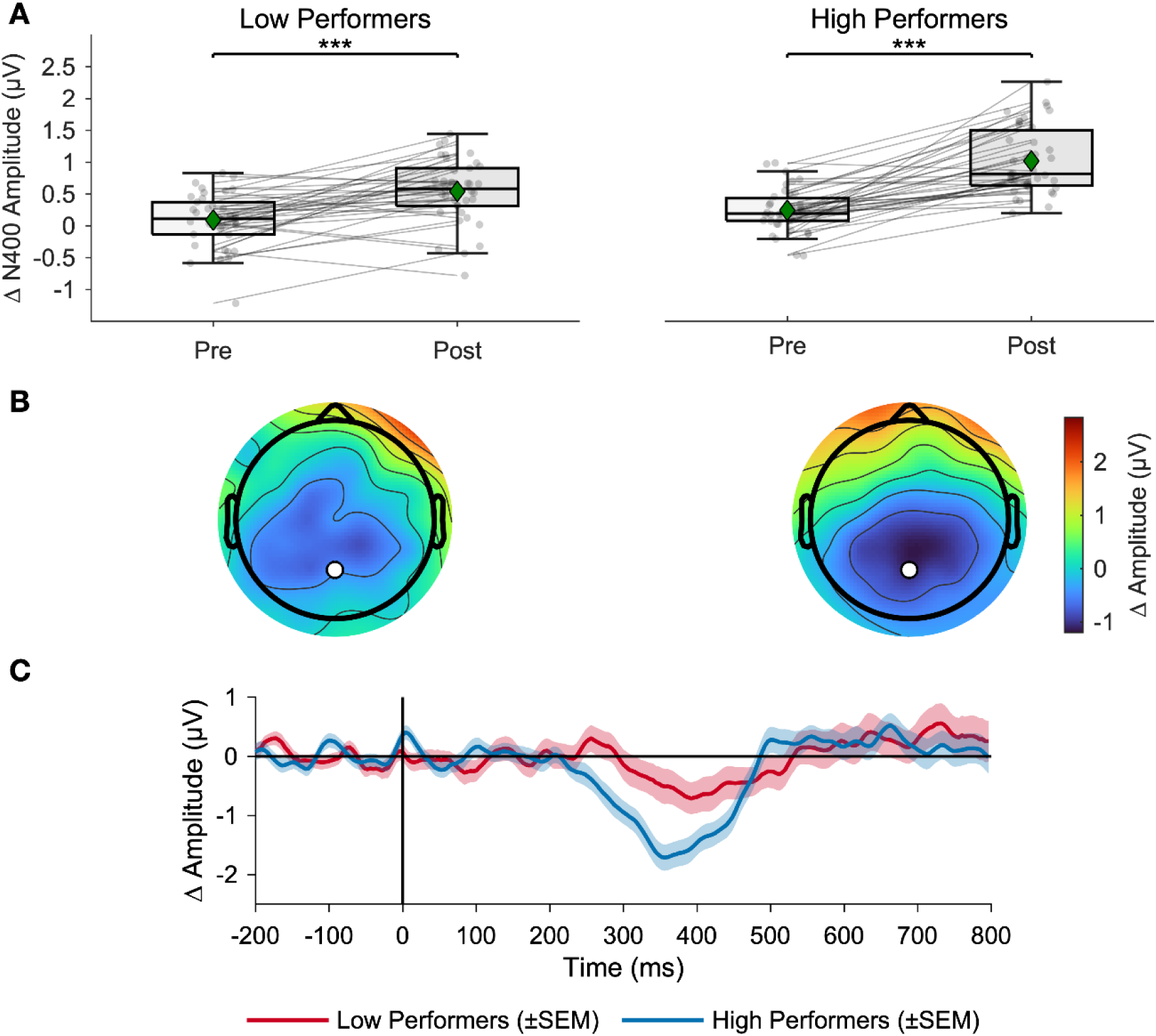
N400 Amplitude Change, Scalp Topographies, and ERP Waveforms. *Note.* Panel A displays individual N400 incongruity effects (incongruent−congruent) before and after learning for low and high performers. Gray lines connect repeated measurements within participants; green diamonds represent group means; asterisks indicate significant within-group differences (*p* < .001). Panel B depicts post–pre scalp topographies of the N400 incongruity effect (300–500 ms), with low performers on the left and high performers on the right; electrode Pz is marked by a white dot (color scale in µV at right). Panel C shows the grand-average post–pre ERP difference wave for the N400 incongruity effect at electrode Pz, with shaded bands representing standard error of the mean.

Separate ANOVAs on post-learning N400 incongruity effects—examining each learning strategy by Performance Group—showed no modulation by strategy (all *p* > .588, for *Supplementary Table S5*). However, small but consistent main effects of Performance Group persisted across strategies (all *p* < .01, *η²_g_* < .13), confirming that high performers maintained larger incongruity effects after learning.

Figure 3B illustrates the scalp distribution of the learning-related change in the N400 incongruity effect, calculated as shown in Equation 6:

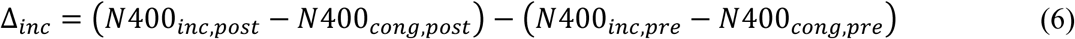

over the 300–500 ms window for low (left) and high (right) performers. Figure 3C presents the grand-average post–pre ERP difference wave at Pz (incongruent−congruent), confirming more pronounced parietal negativity in high performers.

### 3.4 Analyses of Global Features of the Electric Field at the Scalp

#### 3.4.1 Topographic Clustering & GFP

To investigate condition-related changes in the strength and topography of the scalp-recorded electric field, we first examined global features using topographic clustering and Global Field Power (GFP).

Topographic clustering revealed six distinct ERP template maps over the 0–796 ms period (Figure 4A). Across conditions, Global Field Power (Figure 4B) consistently peaked during Map 2 (∼ 148–296 ms), indicating heightened early processing preceding the N400 window. Within the N400 window (300–500 ms), Maps 3 and 4 predominated. Map 3, characterized by broad left-hemispheric negativity, was present in pre-learning congruent and incongruent conditions as well as in the post-learning incongruent condition, but absent in the post-learning congruent condition, which was characterized by Map 4 and uniquely by Map 5. Map 4 displayed right-posterior positivity with opposing left-frontal negativity, while Map 5 featured left-posterior negativity coupled with pronounced right-central positivity.

**Figure 4.**
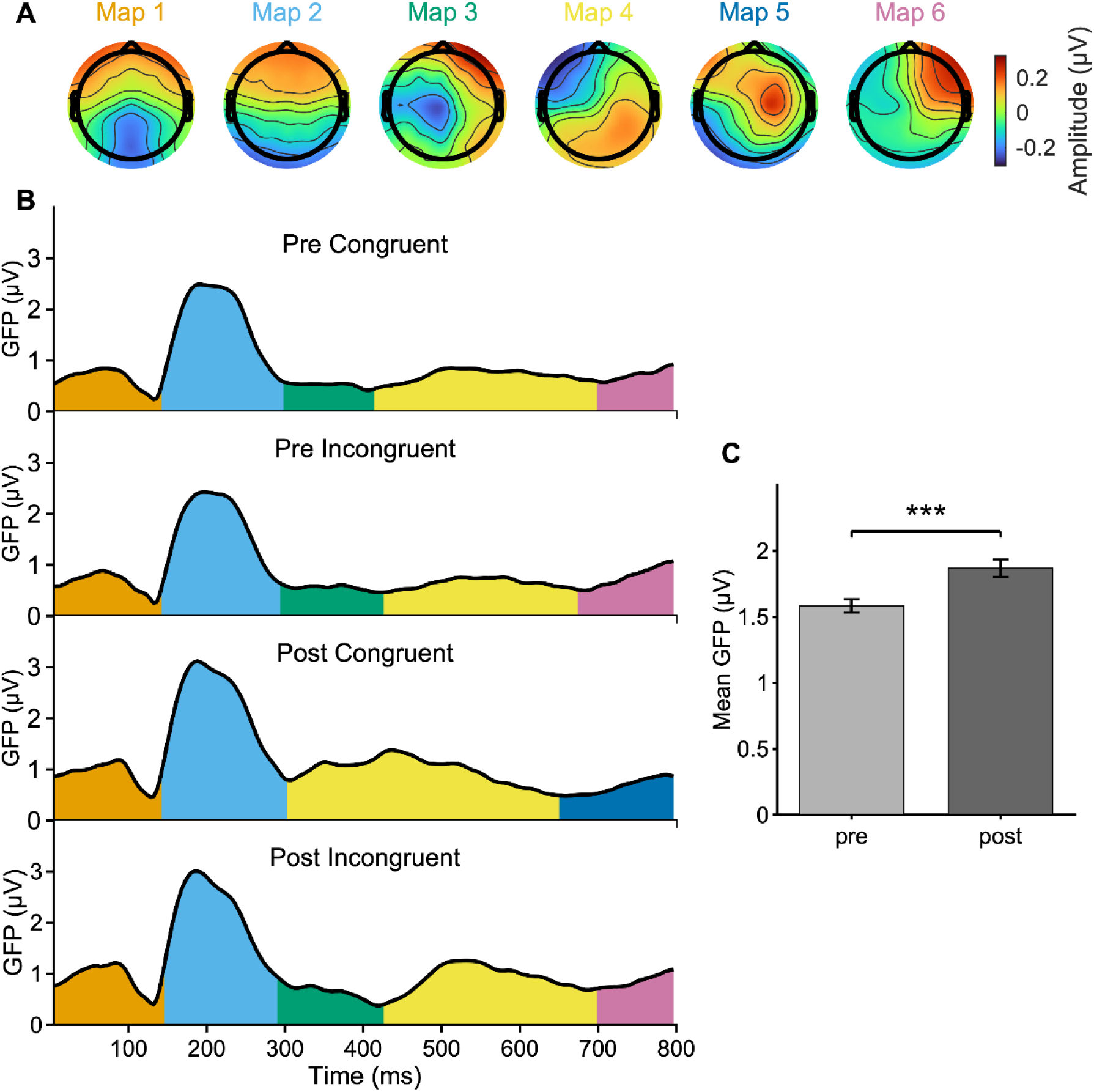
ERP Template Maps, GFP Time Courses by Condition with Segmentation, and Mean GFP by Session. *Note.* Panel A shows the six ERP template maps (µV; color bar at right). Panel B displays GFP time courses for the four experimental conditions, with shaded colored bands indicating the temporal extent of each corresponding map from A. Panel C presents mean GFP amplitudes pre- and post-learning (± *SEM*); paired-samples t-test, *t*(76) = −4.42, *p* < .001.

This pattern suggests a learning-related reorganization of semantic processing. The disappearance of Map 3 post-learning for congruent items likely reflects a reduced need for effortful semantic processing, whereas Map 5’s unique presence suggests a more efficient or specialized processing state.

In line with this interpretation, permutation testing on GFP using the global pAUC statistic revealed a significant main effect of Session (global pAUC, *p* = .0002; Figure 4C), indicating stronger overall brain responses post-learning. No other main effects or interactions were significant (all global pAUC tests, *ps* > .129).

#### 3.4.2 Template-Map Preponderance by Epoch

To investigate how template-map dynamics varied with learning, congruency, and performance, we analyzed the preponderance of each template map within three predefined temporal epochs. Preponderance was defined as the number of time frames (TF) in a participant’s ERP that showed the highest spatial correlation with a given template map, as identified during topographic clustering. Repeated-measures ANOVAs were conducted for each epoch with the factors Map, Session, Congruency, and Performance Group.

Epoch 1 (0–73 TF; 0–292 ms) was dominated by Maps 1 and 2. A main effect of Map, *F*(1, 75) = 10.30, *p* = .002, *η²_g_* = .09, indicated that Map 2 exhibited a higher preponderance than Map 1, independent of Session, Congruency, or Performance Group.

Epoch 2 (74–162 TF; 292–648 ms), which corresponds to the N400 window, was characterized by Maps 3 and 4 (Figure 5 *A*). A Session × Congruency × Map interaction, *F*(1, 75) = 5.43, *p* = .022, indicated condition-specific shifts in semantic processing dynamics. Follow-up analyses showed that, in post-learning congruent trials, Map 4 showed higher preponderance than Map 3, *F*(1, 75) = 22.03, *p* < .001, *η²_g_* = .23, with this effect stronger in high performers (Map × Performance Group: *F*(1, 75) = 6.42, *p* = .013, *η²_g_* = .08).

**Figure 5.**
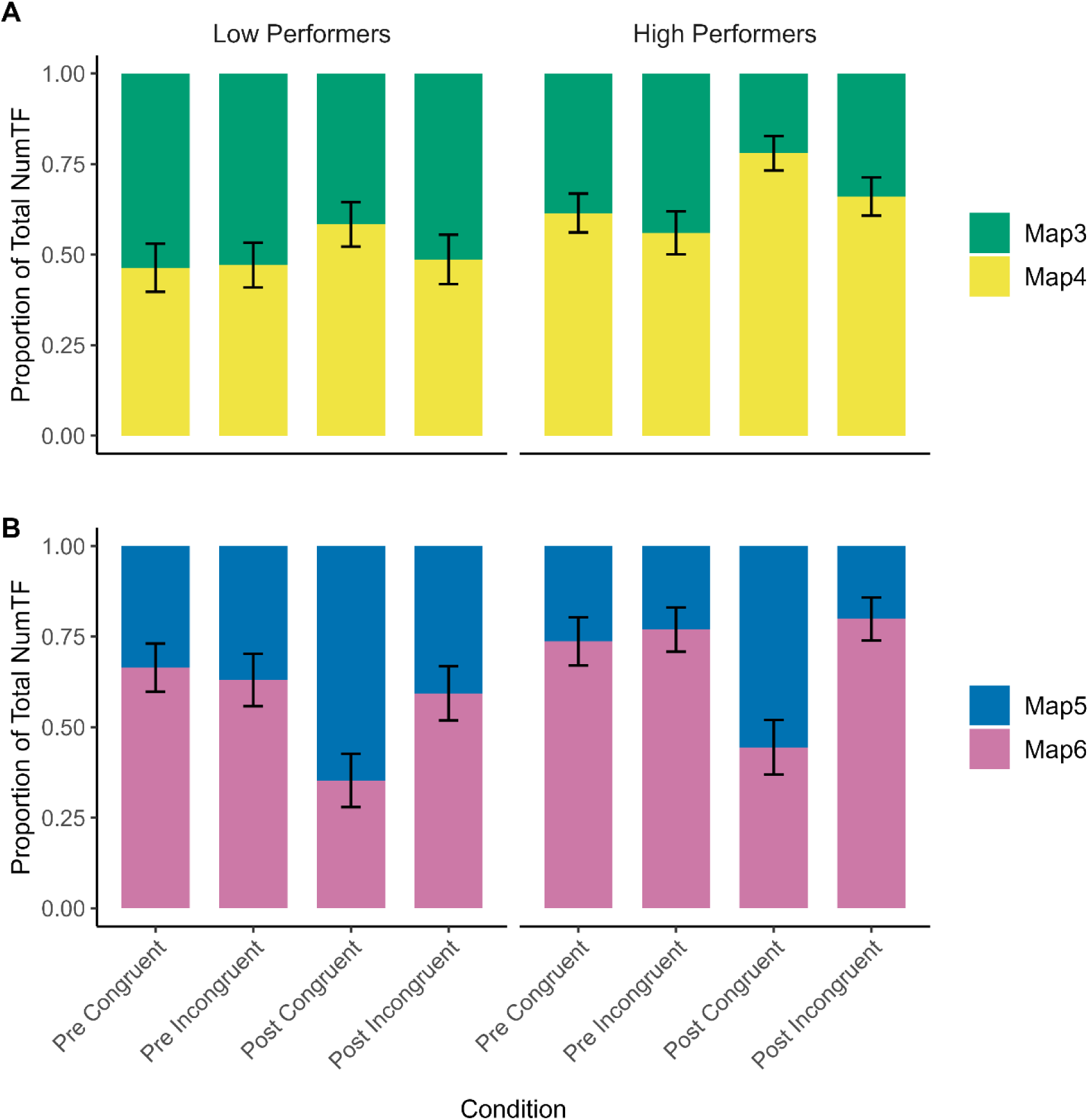
Preponderance of Map 3, 4 and Map 5, 6 by Condition and Performance group. *Note.* Preponderance of Map 3 and 4 (A), and Map 5 and 6 (B) expressed as proportion of total time frames per condition and performance group. Bars represent mean proportions. Error bars indicate the standard error of the mean.

In post-learning incongruent trials, a similar Map × Performance Group trend emerged (*F*(1, 75) = 4.07, *p* = .048, *η²_g_* = .05), although the main effect of Map did not reach significance (*p* = .092). No Map- or Performance-related effects occurred in the pre-learning conditions. Across all conditions, a robust Congruency × Map interaction, *F*(1, 75) = 18.85, *p* < .001, *η²_g_* = .01, showed higher Map 3 preponderance during incongruent trials and higher Map 4 preponderance during congruent trials. A Session × Map interaction, *F*(1, 75) = 8.97, *p* = .004, *η²_g_* = .02, reflected increased Map 4 preponderance after learning.

Epoch 3 (163–199 TF; 648–796 ms) involved Maps 5 and 6 (see Figure 5 *B*). A Session × Congruency × Map interaction, *F*(1, 75) = 19.46, *p* < .001, *η²_g_* = .03), prompted follow-up ANOVAs. In all pre-learning conditions and in the post-learning incongruent condition, Map 6 preponderance exceeded those of Map 5 (all *F* ≥ 16.95, *p* ≤ .001). The post-learning congruent condition showed no significant Map difference (*p* = .058). A Map × Performance Group interaction in the post-learning incongruent condition, *F*(1, 75) = 4.66, *p* = .034, *η²_g_* = .06, indicated greater Map 6 preponderance among high performers. No other Performance Group interactions were observed (*Fs* ≤ 2.17, *ps* ≥ .145). These results suggest that learning effects on late-stage topographic map dynamics were strongest for incongruent items, particularly in high-performing individuals. ANOVA results are provided in *Supplementary Table S6*.

#### 3.4.3 Global Map Dissimilarity

To complement the insights from topographic clustering and GFP, we employed Global Map Dissimilarity (GMD) to quantify, at each time point (0–796 ms), the degree to which the entire scalp topography differed between conditions. Whereas clustering reduces the data to a finite set of recurring ERP template maps and GFP indexes overall field strength, GMD provides a continuous, data-driven metric of spatial configuration differences—even when template-map identity or amplitude remains constant. Permutation testing using the global pAUC statistic revealed significant main effects of Session (global pAUC, *p* < .001), Congruency (global pAUC, *p* < .001), and Performance Group (global pAUC, *p* = .008), as well as significant Session × Congruency (global pAUC, *p* = .010) and Session × Performance Group (global pAUC, *p* = .009) interactions. The TANOVA time windows for the factor Session (the entire period 0–796 ms), for Congruency (from 232 ms onward) and the Session × Congruency interaction (260–476 ms and 508–796 ms; see *Supplementary Figure S1* mirror our clustering and GFP findings.

Notably, the Session × Performance Group interaction was most pronounced between 276 and 428 ms (*Supplementary Figure S1E*), coinciding with the N400 window. This timing aligns with the analysis of topographic map preponderance in Epoch 2, where a Map × Performance Group interaction (*F*(1, 75) = 6.42, *p* = .013, *η²_g_* = .08) showed that high performers exhibited higher Map 4 preponderance than low performers. Thus, the GMD confirms that individual competency not only modulates the strength (GFP) and topographic shifts (clustering) of the N400, but also the continuous scalp-map reconfigurations captured by GMD.

No significant Congruency × Performance Group or the three-way (Session × Congruency × Performance Group) interactions emerged (all *ps* > .26).

## 4 Discussion

The present study demonstrated robust vocabulary learning gains accompanied by clear neural changes. Following two weeks of L2 vocabulary training, participants exhibited significant improvements in vocabulary test scores and enhanced discrimination sensitivity (*d′*) for congruent versus incongruent word pairs. Critically, the N400 incongruity effect, minimal prior to training, emerged prominently after learning, reflecting successful integration of new L2 words into participants’ semantic networks. High-performing learners (those with above-median vocabulary gains) consistently showed larger N400 differences than low performers, indicating more efficient neural processing and deeper semantic encoding. Contrary to our hypotheses, however, there was little evidence that specific learning strategies (retrieval practice, corrective feedback, multisensory learning, or distributed learning) differentially modulated the N400 or behavioral outcomes, except for a modest benefit of corrective feedback on vocabulary test performance. These findings are discussed in the context of the established functional role of the N400 and prior ERP research on L2 vocabulary acquisition, with a particular focus on insights from the ERP template-map analyses.

### 4.1 Neural Sensitivity of the N400

The N400 is a well-established index of semantic integration, lexical access ease, and prediction error (Kuperberg & Jaeger, 2016; Kutas & Federmeier, 2011; Lau et al., 2008; Murray et al., 2024). In line with Hypothesis 1, the pronounced N400 incongruity effect observed after learning confirms that the newly acquired words were efficiently integrated into the semantic network, enabling learners to form stronger lexical-semantic predictions.

Consequently, encountering incongruent word pairs elicited greater semantic processing difficulty due to violated predictions, as indicated by increased N400 amplitudes (Kuperberg & Jaeger, 2016; Lau et al., 2008).

Previous ERP studies of L2 vocabulary acquisition typically employed shorter training intervals—often limited to 3–4 hours—and nonetheless reported detectable, albeit smaller, N400 incongruity effects (e.g., McLaughlin et al., 2004; Pu et al., 2016). Pu et al. (2016) showed that learners exhibited no reliable N400 incongruity effect prior to training but developed a significant, though comparatively moderate, effect after only three hours of instruction, with a clear frontal distribution. Importantly, their study exclusively employed L2–L1 translations, whereas our study tested translation recognition in both L2–L1 and L1– L2 directions, providing a more comprehensive assessment of lexical-semantic integration.

By including both translation directions, our design may not only have promoted more robust and balanced semantic mapping by engaging both forward (L1–L2) and backward (L2–L1) translation processes (see Figure 3 for illustration), but also offered a more ecologically valid model of real-world L2 use. In everyday language contexts, L2 learners frequently encounter and produce words in both directions, and thus this bidirectional paradigm more closely mirrors the challenges and demands of authentic second language communication. According to the Revised Hierarchical Model (Kroll & Stewart, 1994), these directions rely on partly distinct mechanisms—semantic mediation for L1–L2 and more direct lexical links for L2– L1—suggesting that activating both may strengthen cross-linguistic integration. High-performers, in particular, may have benefited from this dual-route activation, leading to stronger N400 incongruity effects.

Similar early frontal N400 patterns have been reported by Yum et al. (2014), who observed enhanced frontal negativity in fast learners after short-term training. Likewise, McLaughlin et al. (2004) demonstrated rapid onset of N400 incongruity effects following very brief passive L2 exposure, suggesting initial stages of lexical acquisition.

In contrast, our extended two-week intervention resulted in substantially stronger behavioral improvements and a more mature neural signature of semantic integration, characterized by clearer centro-parietal distributions and enhanced N400 differentiation between congruent and incongruent word pairs. These results align with findings from Armstrong et al. (2024), who observed a similar maturational shift in N400 topography, moving from fronto-central dominance during early learning toward the canonical centro-parietal distribution following multi-day consolidation. Additionally, Elgort et al. (2014) reported comparable centro-parietal distributions among proficient L2 learners, further supporting the notion that increased lexical proficiency shifts the N400 topography toward canonical patterns.

Moreover, the magnitude of the N400 incongruity effect in the present study varied systematically with participants’ vocabulary gains, supporting Hypothesis 4. This suggests that depth of lexical-semantic knowledge, rather than the specific learning strategies employed, primarily drives neural efficiency**—**that is, the brain’s ability to rapidly and effectively integrate new lexical-semantic information, resulting in more streamlined and differentiated neural responses during language processing**—**and ease of semantic integration. This interpretation is consistent with previous studies emphasizing the relationship between proficiency or learning depth and N400 amplitude modulation (e.g., Elgort et al., 2014; Yum et al., 2014). Qi et al. (2017) further demonstrated that initial N400 amplitudes to native language stimuli predicted subsequent vocabulary and grammar learning in an artificial language, underscoring the role of individual differences in semantic processing efficiency, a concept used interchangeably here with "neural efficiency" for learning outcomes.

### 4.2 Impact of Learning Strategies on Behavioral and Neural Outcomes

One objective was to examine whether established learning strategies would produce measurable differences. Across all participants, a combination of retrieval practice, feedback, multisensory input, and distributed learning resulted in substantial learning gains—approximately a 75% increase in vocabulary scores and improved translation recognition accuracy. However, when analyzing the effects of individual learning strategies, differences were modest, and no strategy reliably modulated the N400 amplitude or behavioral outcomes in isolation.

In line with Hypothesis 2, we expected that retrieval practice would enhance long-term retention and sharpen semantic representations, potentially manifesting as a stronger post-learning N400 incongruity effect or superior behavioral performance. While the overall intervention clearly benefited learning, we did not observe a significant advantage for items practiced with higher retrieval proportion. This stands in contrast to previous findings demonstrating that retrieval practice enhances recall accuracy during vocabulary acquisition—a phenomenon known as the testing effect. Behavioral studies have consistently shown that actively retrieving word associations leads to superior long-term retention compared to passive restudy (Barcroft, 2007; Fritz et al., 2007; Kang et al., 2013; Roediger & Karpicke, 2006). At the neural level, Liu et al. (2018) reported distinct ERP correlates of retrieval-based learning, and Li et al. (2023) found that actively retrieved L3 words elicited a reduced N400 amplitude compared to restudied items, interpreted as reflecting more efficient semantic access. In our study, however, no such effect emerged. This may be due to ceiling effects or overlapping exposure: items were learned repeatedly across conditions, and the high overall success rate may have masked potential condition-specific differences.

A related issue concerns how ERP measures such as the N400 can be meaningfully linked to cognitive processes like encoding richness or memory consolidation. While reduced N400 incongruity is often interpreted as reflecting more fluent lexical-semantic access, these reductions do not necessarily indicate deeper or more stable memory representations. As such, the expected benefits of multisensory input and distributed learning on encoding and retention (Hypotheses 2 and 3) were not reflected in post-learning N400 incongruity or behavioral outcomes. Prior studies have shown that multimodal presentation (e.g., visual and auditory) deepens semantic encoding (Shams & Seitz, 2008), and that spaced repetition enhances memory stability (Cepeda et al., 2006). However, our design did not implement pure “present vs. absent” contrasts for any individual learning strategy. Even "unisensory" items could involve both visual and auditory input, shown in distinct screen regions (e.g., one in the upper and one in the lower part of the display). All participants engaged in repeated practice over a two-week period, which may have diminished the contrasts between conditions. Unlike studies that implemented sharper contrasts—such as multimodal versus unimodal input (e.g., Thelen & Murray, 2013) or massed versus widely spaced repetition at the item level (e.g., Atkin et al., 2023; Feng et al., 2019)—our manipulation may not have been sufficiently strong to yield distinct neural or behavioral effects. Additionally, most of the aforementioned studies used between-subjects designs, which may have amplified contrast effects between learning conditions. Together, these differences in experimental design, contrast strength, and spacing duration may explain why the expected benefits of learning strategies were not observable in our ERP and behavioral outcomes.

Subdividing the sample by learning strategy and performance group may have reduced statistical power to detect small or interaction effects. We did observe a very small behavioral benefit of corrective feedback: across participants, items where the correct answer was immediately presented when a mistake was made were slightly better than those without feedback. This aligns with educational theories that timely feedback strengthens memory traces by explicitly correcting the erroneous associations (Nicol & Macfarlane-Dick, 2006). Yet, the effect size was limited (*η²_g_* ≈ .03), and no corresponding effects emerged at the neural level, such as in the N400 amplitude.

In summary, although the intervention led to substantial gains in vocabulary knowledge and neural differentiation, our results did not provide clear evidence that any individual learning strategy—retrieval practice, corrective feedback, multisensory learning, or distributed learning—consistently enhanced outcomes beyond the general effects of exposure. The small behavioral benefit observed for corrective feedback was specific to low performers—those who showed smaller accuracy gains—suggesting that immediate presentation of the correct answer after errors may be especially helpful for individuals with less improvement in vocabulary retention, even though no corresponding neural effects were found. Given the high overall learning success across conditions, it remains possible that the cumulative exposure across multiple strategies, rather than any specific one, supported lexical-semantic integration. However, since the design did not include a no-strategy control condition, conclusions about additive or synergistic effects of strategy combinations remain speculative. Future studies should more systematically isolate and contrast strategy effects to clarify their unique and interactive contributions to vocabulary learning.

### 4.3 High vs. Low Performers: Neural Efficiency and Strategy Use

High performers showed more pronounced N400 incongruity effects and subtle topographic differences compared to low performers. Our study adds to the literature by showing that even when group-level learning gains are large, ERP measures such as the N400 and ERP template-map analyses reveal persistent qualitative differences between high and low performers—reflecting differences in neural efficiency (as previously defined, i.e., the brain’s ability to rapidly and effectively integrate new lexical-semantic information). In other words, high-performing learners exhibit more efficient neural processing of new vocabulary, as indicated by larger and more consistent N400 incongruity effects and more specialized ERP template-map configurations, which signal deeper semantic integration and more stable memory traces. In contrast, low performers may have remained uncertain about word meanings even post-learning, resulting in reduced or inconsistent N400 responses and less differentiated neural patterns.

This pattern aligns with the idea that proficient learners process L2 semantics more similarly to native speakers (McLaughlin et al., 2004). Our high-performing group’s neural activity can be viewed as more tuned to the semantics of the new words, whereas low performers might rely on more analytical or familiarity-based strategies (potentially engaging later positivity components for decision uncertainty). Although we did not find significant learning strategy-specific improvements for high versus low performers, minor interactions in response bias measures suggest that certain learning strategies (e.g., distributed learning) might subtly modulate response tendencies. Overall, these results highlight ERPs as a sensitive tool for revealing qualitative differences in how deeply L2 words are integrated, beyond simple recall accuracy.

### 4.4 Neural Dynamics Revealed by ERP Template-Map Analyses, GFP, and GMD

By extending traditional ERP analysis with ERP template-map analyses (topographic clustering), GFP, and GMD approaches, we provided a more detailed view of how the spatiotemporal patterns of brain activity are reorganized following vocabulary learning. This multimethod approach provided a more fine-grained view of when and how the brain’s functional states differ before versus after vocabulary acquisition.

We identified six distinct ERP template maps that describe the dominant EEG topographies across the 800 ms epoch. Within the canonical N400 window (300–500 ms), we observed two different ERP template maps (Maps 3 and 4) whose preponderance varied by condition—a finding in line with Brandeis et al. (1995), who reported separate preN400 and N400 scalp-maps during sentence processing. Notably, Map 3 (left-hemispheric negativity) was prominent before learning and in post-learning incongruent trials, while Map 4 and a unique Map 5 (right-central positivity/left-posterior negativity) characterized post-learning congruent trials, suggesting a shift towards more fluent or automatized processing. High performers exhibited higher Map 3 preponderance within the N400 window, consistent with their larger N400 amplitudes, indicating sustained semantic processing.

This finding—that the post-learning congruent condition occupies a unique neural state—aligns with the emergence of Map 5 and underscores a qualitative change in how known correct translations are processed. Comparable shifts in scalp topographies driven by semantic predictability have also been reported in sentence processing (e.g., Perrin & Garcia-Larrea, 2003) and in category learning and semantic decision tasks (Khateb et al., 2003), suggesting that semantic congruency can induce distinct neural configurations. Furthermore, findings from auditory learning (Spierer et al., 2007) and perceptual classification (Tzovara et al., 2012) provide theoretical and empirical support for the idea that learning systematically reorganizes not only response amplitude but also the spatial dynamics of brain activity across domains.

Our GFP analysis revealed a general increase in global field strength post-learning— regardless of congruency—implying increased overall neural engagement.

GMD confirmed robust congruency effects beginning at 232 ms, consistent with the onset of semantic processing reported in the literature (Hauk et al., 2006; Kutas & Federmeier, 2011; Lau et al., 2008). The overall topographic pattern of the Session × Congruency interaction mirrors the learning- and congruency-related reorganization observed in the clustering and GFP results, reflecting conceptual convergence across analytic methods without offering additional temporal detail.

In sum, these results demonstrate that vocabulary learning enhances both the strength and the spatiotemporal configuration of neural activity during semantic processing, as measured by multiple complementary EEG analysis techniques.

### 4.5 Comparison with Previous Studies and Implications

Our findings contribute to a growing body of research at the intersection of L2 learning and neurocognition. They corroborate prior evidence that even initial stages of L2 word learning modulate the N400 (Pu et al., 2016; Kutas & Federmeier, 2011) and extend this by linking the magnitude of the N400 incongruity effect to individual vocabulary gains (Qi et al., 2017; Elgort et al., 2014).

Although we tested four evidence-based digital learning strategies—retrieval practice, corrective feedback, multisensory encoding, and distributed learning—none showed clear advantages over the others in our design. This absence of differential strategy effects mirrors well-known boundary conditions: Roediger and Karpicke (2006) demonstrated that, once recall approaches ceiling levels, additional retrieval practice yields diminishing gains; and Kornell and Bjork (2007) showed that when retrieval is too easy (i.e., items are already well learned), testing produces smaller benefits than under more challenging conditions. Together, these studies indicate that when learning items are overlearned or tasks lack sufficient difficulty, the unique contributions of individual strategies are masked—exactly as we observed after our high-exposure, high-success learning intervention.

Beyond educational implications, our analysis of topographic ERP mapping illuminates how vocabulary learning is accompanied by neural reorganization. The clear distinction between processing known versus unknown words supports models of synaptic and systems consolidation, whereby repeated activation refines synaptic weights and engages hippocampal–neocortical interactions to stabilize memory traces over time (McGaugh, 2000; Frankland & Bontempi, 2005). This “representational sharpening” emerges as reduced trial-to-trial variability and the appearance of specialized EEG map configurations (Paller & Wagner, 2002; Yassa & Stark, 2011). Consistent with the neural efficiency accounts—which posit that increasing proficiency yields more selective and topographically focal neural engagement (Haier et al., 1992; Neubauer & Fink, 2009)—our topographic ERP mapping findings show increasingly focal scalp topographies both after vocabulary acquisition and in high-performing learners (Khateb et al., 2003; Spierer et al., 2007; Tzovara et al., 2012).

Importantly, whereas Neubauer and Fink (2009) report reduced global field strength in experts, our high performers exhibited larger N400 incongruity effects alongside focal ERP template-map engagement, reflecting qualitative specialization rather than mere amplitude suppression. Together, these results provide empirical support for theories of brain plasticity showing that, as vocabulary knowledge consolidates, cortical networks reorganize into more stable, specialized patterns observable across perceptual, semantic, and associative learning domains.

These findings underscore the potential of ERP and topographic clustering as sensitive tools for monitoring individual progress and for evaluating the impact of pedagogical interventions in L2 learning.

Future research should investigate whether these learning-induced neural signatures— such as changes in N400 incongruity and ERP template-map dynamics—generalize to other populations and tasks, including productive language use. In particular, children typically show shorter and less stable EEG microstates in resting activity (Koenig et al., 2002) and delayed/broader N400 (Holcomb et al., 1992; Friedrich & Friederici, 2004).

### 4.6 Limitations

There are some limitations to consider. First, although we manipulated three strategies (retrieval practice, corrective feedback, multisensory learning) within participants and distributed learning between participants, the nested implementation may have attenuated contrasts. Moreover, the two-week intervention yielded very high overall success rates— vocabulary test performance increased from 0.41% to 75.5% (*d_z_* = 3.96) and sensitivity (*d*′) in the translation recognition task increased from 0.59 to 4.15 (*η²_g_* = .89)—such that even sub- optimal strategy assignments were sufficient to drive near-ceiling learning and obscured any strategy-specific benefits (all *η²_g_* ≤ .03 for strategy factors). Second, although our EEG sample (N = 77) is large for the field, exclusions for artifact rejection and the median split reduced power for interaction tests; effect sizes for strategy-related ERP interactions were small (*η²_g_* ≤ .02). Third, our EEG paradigm focused exclusively on translation-recognition— a receptive task in both L2–L1 and L1–L2 directions—and did not assess productive use (e.g., cued recall or sentence production). It remains to be tested whether similar ERP template-map reconfigurations and N400 dynamics arise under more generative demands, which would help establish the generality of these neural markers across task contexts.

### 4.7 Conclusion

This study demonstrates that intensive L2 vocabulary learning rapidly strengthens semantic processing, as evidenced by larger N400 incongruity effects in successful learners, while remaining largely unaffected by the particular learning strategy employed. Our ERP template-map analyses further demonstrate that learning not only shifts which ERP template maps dominate the N400 window from pre- to post-learning, but also differentially by congruency (congruent vs. incongruent), reflecting a qualitative reorganization of neural activity rather than a mere increase in amplitude. Together, these results bridge cognitive neuroscience and language education by demonstrating that effective vocabulary acquisition leaves stable, condition-specific neural signatures in semantic processing.

Importantly, our findings underscore the practical value of ERP and topographic clustering for capturing individual learning-related changes in second language acquisition— even in the absence of clear differential effects of specific learning strategies. These neural measures may therefore provide sensitive biomarkers of L2 learning success, offering unique insights into individual differences in learning outcomes that are not always captured by behavioral measures alone.

## Author Contributions

Nicole H. Skieresz: Conceptualization; Methodology; Investigation; Formal analysis; Data curation; Writing – original draft; Writing – review & editing.

Sandy C. Marca: Investigation; Data curation; Methodology; Writing – review & editing.

Micah M. Murray: Supervision; Conceptualization; Methodology; Formal analysis; Writing – review & editing.

Thomas P. Reber: Supervision; Conceptualization; Methodology; Writing – review & editing; Funding acquisition.

Nicolas Rothen: Supervision; Conceptualization; Methodology; Writing – review & editing; Funding acquisition.

*Note.* Micah M. Murray, Thomas P. Reber and Nicolas Rothen contributed equally to this work (*).

## Data Availability Statement

Data and analysis scripts are available to editors and reviewers via a private, view-only OSF link: https://osf.io/bj6pt/?view_only=ead3461634134e209cd8c234687e1d6e

Upon acceptance, we will make the OSF project public. Preprocessed EEG data are archived on Zenodo (DOI: https://doi.org/10.5281/zenodo.16538184) and are currently accessible to reviewers via a private link described in the OSF README. The Zenodo record will be made public upon publication.

