## Supplementary Material for "Brain Network Differences in Second Language Learning Depend on Individual Competencies"

**Table S1**

*Full Stimulus Characteristics for All Items*

| German (syll./len./Zipf) | French (syll./len./Zipf) | Finnish (syll./len./Zipf) | English (translation) |
|---|---|---|---|
| Vogel (2/5/4.45) | oiseau (2/6/4.31) | lintu (2/5/4.31) | bird |
| Hund (1/4/4.86) | chien (2/5/4.86) | koira (2/5/5.04) | dog |
| Katze (2/5/4.49) | chat (1/4/4.76) | kissa (2/5/4.6) | cat |
| Wolf (1/4/4.57) | loup (1/4/4.36) | susi (2/4/4.23) | wolf |
| Esel (2/4/3.91) | âne (2/3/3.8) | aasi (2/4/3.67) | donkey |
| Kuh (1/3/4.04) | vache (2/5/4.14) | lehmä (2/5/4.04) | cow |
| Schaf (1/5/3.73) | mouton (2/6/3.92) | lammas (2/6/3.85) | sheep |
| Hirsch (1/6/3.85) | cerf (1/4/3.78) | peura (2/5/3.6) | deer |
| Hahn (1/4/4.29) | coq (1/3/3.83) | kukko (2/5/3.78) | rooster |
| Wal (1/3/3.46) | baleine (3/7/3.73) | valas (2/5/3.26) | whale |
| Affe (2/4/3.72) | singe (2/5/3.93) | apina (3/5/3.83) | monkey |
| Rabe (2/4/3.42) | corbeau (2/7/3.49) | korppi (2/6/3.45) | raven |
| Schlange (2/8/4.11) | serpent (2/7/4.02) | käärme (3/6/3.71) | snake |
| Fliege (2/6/3.81) | mouche (2/6/3.9) | lentää (2/6/4.62) † | fly |
| Ente (2/4/3.78) | canard (2/6/4.17) | ankka (2/5/4.07) | duck |
| Donner (2/6/3.57) | tonnerre* (3/6/3.81) | ukkonen (3/7/3.57) | thunder |
| Regen (2/5/4.63) | pluie (2/5/4.56) | sade (2/4/4.16) | rain |
| Wind (1/4/4.69) | vent (1/4/4.8) | tuuli (3/5/4.57) | wind |
| Fluss (1/5/4.44) | fleuve (2/6/4.32) | joki (2/4/4.01) | river |
| Schnee (1/6/4.45) | neige (2/5/4.56) | lumi (2/4/4.17) | snow |
| Tropfen (2/7/4.15) | goutte (2/6/4.03) | kihti (2/5/2.7) † | drop |
| Adler (2/5/4.19) | aigle (2/5/3.82) | kotka (2/5/4.28) | eagle |
| Eule (2/4/3.5) | hibou (2/5/3.23) | pöllö (2/5/3.36) | owl |
| Henne (2/5/3.38) | poule (2/5/4.21) | kana (2/4/4.09) | hen |
| Boot (1/4/4.52) | bateau (2/6/4.61) | vene (2/4/4.23) | boat |
| Füller (2/6/3.09) | stylo (2/5/3.85) | kynä (2/4/3.89) | pen |
| Ball (1/4/4.64) | balle (2/5/4.6) | pallo (2/5/4.63) | ball |
| Dose (2/4/3.91) | canette* (2/7/3.21) | tölkit (2/6/2.8) † | can |
| Hammer (2/6/4.4) | marteau (2/7/3.87) | vasara (3/6/3.75) | hammer |
| Schere (2/6/3.78) | ciseaux (2/7/3.54) | sakset (3/6/3.57) | scissors |
| Tafel (2/5/4.17) | tableau (2/7/4.82) | lauta (3/5/3.51) † | board |
| Tür (1/3/4.9) | porte (2/5/5.47) | ovi (2/3/4.48) | door |
| Flugzeug (2/8/4.46) | avion (2/5/4.86) | kone (2/4/4.84) † | airplane |
| Foto (2/4/5.02) | photo (2/5/5.14) | kuva (2/4/5.43) | photo |
| Korken (2/6/3.01) | bouchon (2/7/3.79) | pistoke (3/7/2.51) † | cork |
| Rakete (3/6/3.89) | missile (3/7/3.84) | ohjus (3/5/3.79) | rocket |

| **German (syll./len./Zipf)** | **French (syll./len./Zipf)** | **Finnish (syll./len./Zipf)** | **English (translation)** |
|---|---|---|---|
| Feder (2/5/3.91) | ressort (3/7/4.25) | kevät (2/5/4.43) † | spring |
| Säge (2/4/3.37) | scie (1/4/3.62) | näin (2/4/6.04) † | saw |
| Pfeife (2/6/3.6) | sifflet (2/7/3.39) | pilli (2/5/3.38) | whistle |
| Trommel (2/7/3.55) | tambour (3/7/3.65) | rumpu (2/5/3.23) | drum |
| Hupe (2/4/3.09) | klaxon (2/6/3.01) | tuutata (3/7/1.53) † | horn |
| Teller (2/6/4.09) | assiette (3/8/4.07) | lautanen (3/8/3.44) | plate |
| Schwert (1/7/4.1) | épée (2/4/4.2) | miekka (3/6/3.84) | sword |
| Kanone (3/6/3.44) | canon (2/5/4.36) | tykki (2/5/3.7) | cannon |
| Besen (2/5/3.49) | balai* (2/6/3.36) | luuta (2/5/3.48) | broom |
| Geld (1/4/5.59) | argent (2/6/5.37) | raha (2/4/4.86) | money |
| Flöte (2/5/3.54) | flute (2/5/3.65) | huilu* (2/5/3.15) | flute |

*Note*. Syll. = number of syllables; len. = length in letters; Zipf = Zipf frequency. English translations are provided for illustration only; only the original items were used as experimental stimuli. Words marked with an asterisk (*) contained typos and were excluded from analyses. Words marked with a dagger (†) are Finnish translation errors (i.e., the word used in the experiment did not match the intended meaning); these items were included in the experiment; see Methods.

**Table S2**

*Vocabulary Test: Effects of Learning Strategies on Post-Learning Scores*

| **Predictor** | ***F(df1, df2)*** | ***p*** | $\eta^2_g$ |
|---|---|---|---|
| Performance Group | > 86.80 (1, 79–81) | < .001 | 0.45–0.56 |
| Retrieval Practice | 0.06 (1, 81) | .807 | 0.00 |
| Corrective Feedback | 10.95 (1, 80) | .001 | 0.03 |
| Multisensory Learning (Encoding) | 0.81 (1, 81) | .372 | 0.00 |
| Multisensory Learning (Retrieval) | 0.01 (1, 81) | .905 | 0.00 |
| Distributed Learning | 2.53 (1, 79) | .116 | 0.03 |

*Note.* $F(1, 81)$ refers to the F-statistic with degrees of freedom (*df1, df2*). *df1* indicates degrees of freedom numerator. *df2* indicates degrees of freedom denominator. $p$ = significance level. $\eta^2_g$ = generalized eta squared, indicating effect size.

**Table S3**

*Effects of Performance Group and Session on d' in the Translation Recognition Task*

| **Predictor** | ***F(df1, df2)*** | ***p*** | $\eta^2_g$ |
|---|---|---|---|
| Session | 1603.27 (1, 75) | < .001 | .89 |
| Performance Group | 30.16 (1, 75) | < .001 | .20 |
| Session × Performance Group | 37.51 (1, 75) | < .001 | .16 |
| Retrieval Practice | 0.00 (1, 75) | .988 | .00 |
| Corrective Feedback | 3.16 (1, 74) | .080 | .01 |
| Multisensory Learning (Encoding) | 0.03 (1, 75) | .874 | .00 |
| Multisensory Learning (Retrieval) | 0.36 (1, 75) | .551 | .00 |
| Distributed Learning | 0.12 (1, 73) | .727 | .00 |

*Note.* $F(1, 75)$ refers to the F-statistic with degrees of freedom *(df1, df2)*. *df1* indicates degrees of freedom numerator. *df2* indicates degrees of freedom denominator. $p$ values are two-tailed. $\eta^2_g$ = generalized eta squared effect size measure.

**Table S4**

*Effects of Performance Group and Session on Criterion in the Translation Recognition Task*

| **Predictor** | ***F(df1, df2)*** | ***p*** | $\eta^2_g$ |
|---|---|---|---|
| **Performance Group** | 5.92 (1, 75) | .017 | .04 |
| **Session** | 9.49 (1, 75) | .003 | .05 |
| Performance Group × Session | 0.75 (1, 75) | .390 | .00 |
| Retrieval Practice (RP) | 0.07 (1, 75) | .788 | .00 |
| Performance Group × RP | 0.07 (1, 75) | .794 | .00 |
| Corrective Feedback | 1.26 (1, 74) | .265 | .00 |
| Performance Group × CF | 0.29 (1, 74) | .590 | .00 |
| Multisensory Learning at Encoding (EM) | 0.59 (1, 75) | .443 | .00 |
| Performance Group × EM | 0.25 (1, 75) | .620 | .00 |
| Multisensory Learning at Retrieval (RM) | 1.76 (1, 75) | .189 | .00 |
| **Performance Group × RM** | 4.67 (1, 75) | .034 | .01 |
| Distributed Learning (DL) | 0.40 (1, 73) | .528 | .01 |
| **Performance Group × DL** | 5.40 (1, 73) | .023 | .07 |

*Note***.** *F*(1, 75) refers to the F-statistic with degrees of freedom *(df1, df2)*. *df1* indicates degrees of freedom numerator. *df2* indicates degrees of freedom denominator. *p* values are two-tailed. $\eta^2_g$ = generalized eta squared effect size measure. Bold highlighted predictors represent significant effects.

**Table S5**

*N400 Amplitude Difference (Incongruent Minus Congruent) by Performance Group, Session, and Learning Strategy*

| **Model** | **Predictor** | ***F(df1, df2)*** | ***p*** | $\eta^2_g$ |
|---|---|---|---|---|
| **Main Analysis** | Performance Group | 14.44 (1, 75) | .000 | 11 |
| | Session | 99.52 (1, 75) | .000 | .32 |
| | Performance Group × Session | 6.79 (1, 75) | .011 | .03 |
| **Retrieval Practice** | Performance Group | 19.73 (1, 75) | .000 | .13 |
| | Retrieval Practice (RP) | 0.00 (1, 75) | .978 | .00 |
| | Performance Group × RP | 0.74 (1, 75) | .394 | .00 |
| **Corrective Feedback** | Performance Group | 12.40 (1, 74) | .001 | .08 |
| | Corrective Feedback (CF) | 0.01 (1, 74) | .927 | .00 |
| | Performance Group × CF | 0.15 (1, 74) | .703 | .00 |
| **Multisensory Learning (Encoding)** | Performance Group | 18.86 (1, 75) | .000 | .14 |
| | Encoding (EM) | 0.03 (1, 75) | .861 | .00 |
| | Performance Group × EM | 3.65 (1, 75) | .060 | .02 |
| **Multisensory Learning (Retrieval)** | Performance Group | 17.12 (1, 75) | .000 | .13 |
| | Retrieval (RM) | 0.00 (1, 75) | .961 | .00 |
| | Performance Group × RM | 0.00 (1, 75) | .967 | .00 |
| **Distributed Learning** | Performance Group | 15.81 (1, 73) | .000 | .18 |
| | Distributed Learning (DL) | 0.30 (1, 73) | .588 | .00 |
| | Performance Group × DL | 0.11 (1, 73) | .745 | .00 |

*Note.* All models by learning strategies tested post-learning N400 differences (incongruent – congruent). $F(1, 75)$ refers to the F-statistic with degrees of freedom *(df1, df2)*. *df1* indicates degrees of freedom numerator. *df2* indicates degrees of freedom denominator. *p* values are two-tailed. $\eta^2_g$ = generalized eta squared effect size measure.

**Table S6**

*Topographic Map Preponderance by Map, Session, Congruency, and Performance Group*

| **Model** | **Predictor** | ***F(df1, df2)*** | ***p*** | $\eta^2_g$ |
|---|---|---|---|---|
| **Epoch 1 (TF 1–73)** | Map | 10.30 (1, 75) | .002 | .09 |
| **Epoch 2 (TF 74–163)** | Map | 4.53 (1, 75) | .037 | .04 |
| | Performance Group × Map | 4.38 (1, 75) | .040 | .04 |
| | Session × Map | 8.97 (1, 75) | .004 | .02 |
| | Congruency × Map | 18.85 (1, 75) | < .001 | .01 |
| | Session × Congruency × Map | 5.43 (1, 75) | .022 | .00 |
| **Epoch 3 (TF 164–199)** | Map | 12.79 (1, 75) | .001 | .08 |
| | Performance Group × Map | 3.38 (1, 75) | .07 | .02 |
| | Session × Map | 11.19 (1, 75) | .001 | .03 |
| | Congruency × Map | 14.83 (1, 75) | < .001 | .03 |
| | Session × Congruency × Map | 19.46 (1, 75) | < .001 | .03 |

*Note.* *F*(1, 75) refers to the F-statistic with degrees of freedom *(df1, df2)*. *df1* indicates degrees of freedom numerator. *df2* indicates degrees of freedom denominator. *p* values are two-tailed. $\eta^2_g$ = generalized eta squared effect size measure.

**Figure S1**

*Permutation p value Trajectories Over Time for Significant Global pAUC Tests*

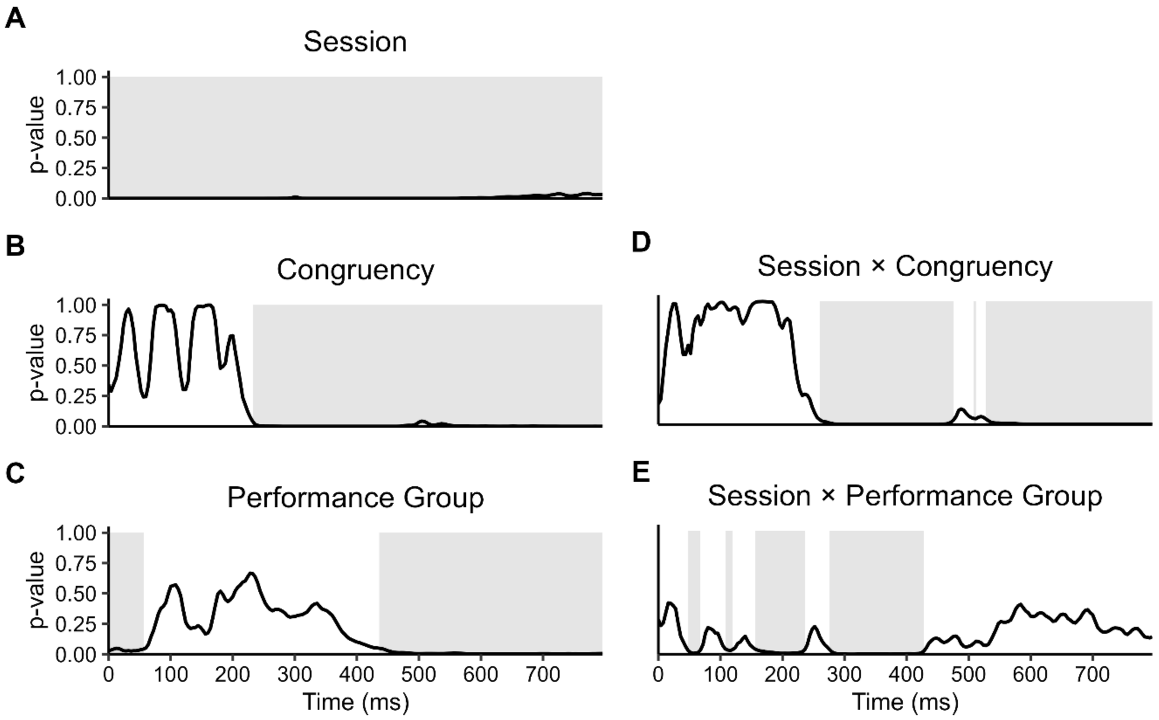


*Note.* Each panel (A–E) shows the permutation *p* value over time for the corresponding global pAUC test across the 0–796 ms epoch. Panels A–C (left column) display main effects of Session, Congruency, and Performance Group; panels D–E (right column) display the Session × Congruency and Session × Performance Group interactions. Gray shading indicates time windows with $p < .05$.
